# A microbial perspective on the life-history evolution of marine invertebrate larvae: if, where, and when to feed

**DOI:** 10.1101/210989

**Authors:** Tyler J. Carrier, Jason Macrander, Adam M. Reitzel

## Abstract

The feeding environment for planktotrophic larvae has a major impact on development and progression towards competency for metamorphosis. High phytoplankton environments that promote growth often have a greater microbial load and incidence of pathogenic microbes, while areas with lower food availability have a lower number of potential pathogens. Trade-offs between metabolic processes associated with growth and immune functionality have been described throughout the animal kingdom and may influence the life-history evolution of marine invertebrate planktotrophic larvae in these environments. Namely, to avoid potential incidences of microbial-mediated mortality and/or dysbiosis, larvae should regulate time spent between these two feeding environments. We describe here transcriptomic and microbiome data that supports this trade-off in larvae, where larvae in a well-fed environment upregulate genes associated with metabolism and may regularly enter a state of dysbiosis, resulting in mortality. To address the hypothesis that the environmental microbiota is a selective force on if, where, and when planktotrophic larvae should feed, we present a strategy for determining the specific interactions of larvae and microbes at a scale representative of their larger pelagic environment.

---

> *“Life in the sea cannot be understood without understanding the sea itself” – Alfred C. Redfield*

## Oceanography and larval evolution

The evolution of life-histories in the sea has largely been shaped by a diverse and interacting suite of oceanographic features (Strathmann 1990; Burgess, Baskett et al. 2015). The oceanographic environment can be divided into four tiers of potential selective pressures: physical, chemical, biological, and microbial. Among these, the first three – physical, chemical, and biological – have been the focus of a majority of life-history research, especially in studying the evolutionary ecology of marine invertebrate larvae (Carrier, Reitzel et al. 2017). For example, the sensory systems of larvae encompassing disparate phyla with distant phylogenetic relationships use physical features of the sea spanning magnitudes in spatial scale, such as sound, turbulence, and olfactory, for navigation (Hodin, Ferner et al. 2017). Additionally, some sea urchin larvae (*e.g.*, *Strongylocentrotus purpuratus* along the Northeastern Pacific Ocean) are capable of exhibiting signs of genomic and physiological adaptions to local ambient acidity and resistance to acidification (Pespeni, Sanford et al. 2013). Lastly, some echinod and mollusc larvae are polyphenic, whereby the biological oceanographic regime, namely high or low levels of phytoplankton, selects for the expression of phenotypes geared toward better feeding performance in each regime (McAlister and Miner 2017).

Pathogenic microbes are known to influence the survival of numerous marine species, particularly in coastal species exposed to anthropogenic encroachment as well as ongoing climate change (Harvell, Mitchell et al. 2002). To date, microbial oceanography has been given little attention when linking the microbial oceanographic environment with life-histories of marine invertebrate larvae. However, recent efforts suggest that the microbiome of marine invertebrate larvae is species-specific and distinct from the environmental microbiota (Galac, Bosch et al. 2016), which is consistent with other animals and their developmental stages (*e.g.*, McFall-Ngai and Ruby 2000). The composition and structure of this associated microbial community is also, in part, influenced by the abiotic and biotic environment experienced by the larva (Webster, Botte et al. 2011; Carrier and Reitzel In Review). The ways by which the environmental microbiota influences the structure and composition of the host-associated microbiome remains largely unexplored, which may be a significant omission in understanding larval life histories because associated microbiota have important impacts on physiology, resistance to pathogens, development, and stress tolerance (Rosenberg, Sharon et al. 2009; McFall-Ngai, Hadfield et al. 2013; Bordenstein and Theis 2015; Gilbert, Bosch et al. 2015; Theis, Dheilly et al. 2016).

Here, we propose the hypothesis that the environmental microbiota is a selective force on if, where, and when planktotrophic larvae of benthic marine invertebrates should feed. We specifically emphasize roles for both pathogenic bacteria and a resultant state of dysbiosis. First, we discuss when and where larvae are likely to interact with pathogenic microbiota in the environment and, by analyzing transcriptomic data for one urchin species, show for the first time the trade-off of metabolic activity and immunity in a marine invertebrate larva. This is followed by a discussion on the influence of the environmental microbiota on the state of the hologenome, which we support with sequenced-based analyses of the microbiota associated with healthy and dying echinoid larvae. Finally, we conclude with a sampling strategy for determining the specific interactions of dilute densities of larvae and variable concentrations of microbes in their natural environments.

## Larval oceanography and the plankton community

Benthic marine invertebrates with planktotrophic larvae typically release their gametes in synchronicity with the initiation of the spring phytoplankton bloom, as a means to maximize the period of high food availability for developing larvae (Starr, Himmelman et al. 1990). Over the course of their planktonic period, planktotrophic larvae experience structural shifts in the phytoplankton community, as it varies dynamically in space (*e.g.*, distance from shore, alongshore, and with depth), time (*e.g.*, daily and seasonally), and diversity (*e.g.*, community members). On a finer scale, the most dominant contributor of the phytoplankton community differs on a per day basis over the course of a bloom (Needham and Fuhrman 2016). During this time, archaeal, bacterial, and likely viral communities exhibit similar daily succession patterns (Needham and Fuhrman 2016), implying that archaeal-bacterial-phytoplankton daily successional patterns are biologically coupled and subsequently contribute to the rapid microbial growth and turn-over during this period (Needham and Fuhrman 2016).

Phytoplankton blooms are tightly regulated by the environmental microbiota (Azam 1998; Needham and Fuhrman 2016). During phytoplankton blooms, dissolved (*e.g.*, dissolved organic material, DOM) and particulate nutrients (*e.g.*, phytoplankton) are two of the primary energy sources for bacteria. Additionally, daily archaeal-bacterial-phytoplankton succession mediate increases in bloom-associated environmental microbial load and DOM (McKenny and Allison 1995; Azam and Malfatti 2007). Some of the environmental microbiota (specifically bacteria and bacteriophages) colonize phytoplankton cells, while others take up and metabolize DOM (Brum, Ignacio-Espinoza et al. 2015; Sunagawa, Coelho et al. 2015). Moreover, in high nutrient environments favoring growth, some bacteria express virulence factors and other pathogenic characteristics (*e.g.*, McKenny and Allison 1995).

In addition to concentrating phytoplankton during feeding, planktotrophic larvae also encounter environmental microbiota. For example, echinopleuti are predicted to contact as many as ~2.0 × 10^7^ bacteria●day^−1^ by feeding alone (Hart and Strathmann 1994; Azam and Malfatti 2007), a portion of which would be consumed directly (Jaeckle 2017). However, this estimate represents a fraction of the total environmental microbiota encountered daily, as bacteria may contact other surfaces of the echinopluteus (McEdward 1984). Thus, it is inevitable that larvae interact with and ingest phytoplankton bloom-associated microbes (*e.g.*, Rivkin, Bosch et al. 1986; Gallagher, Waterbury et al. 1994), making it more likely for larvae to encounter pathogenic or virulent species of microbes (*e.g.*, McKenny and Allison 1995; Azam and Malfatti 2007). Phytoplankton, particularly during a spring bloom, can be concentrated and provided enhanced levels of both light and nutrients for growth at two common physical oceanographic phenomena: pycnoclines and frontal zones. Planktotrophic larvae, as well as phytoplankton communities, tend to aggregate in areas directly adjacent to these physical oceanographic features (Dekshenieks, Donaghay et al. 2001; Metaxas, Mullineaux et al. 2009). This is, in part, because larvae are capable of positioning themselves vertically in the water column despite being poor swimmers (Strathmann and Grunbaum 2006; Metaxas, Mullineaux et al. 2009; Arellano, Reitzel et al. 2012). Physical mechanisms and/or behaviors that result in dense aggregations of phytoplankton and associated environmental microbiota may affect life-history trade-offs for larvae. By inhabiting zones that exhibit oceanographic characteristics favoring higher amounts of food, larvae may promote growth and developmental progression. Conversely, these high productivity areas may increase the incidence of interacting with potentially pathogenic microbiota, a potentially significant source of larval mortality (Young and Chia 1987; Rumrill 1990).

To avoid inhabiting positions in the water column where microbe-induced mortality would be higher, larvae may vertically position themselves in zones of reduced phytoplankton abundance and bloom-associated microbiota, including surface waters (incidence of photo-inhibition) and bottom waters (below the critical depth). In these areas microbe-induced mortality may be reduced, while the combined abundance of phytoplankton and alternative dietary options remain sufficient to maintain the larval structures or developmental progression, albeit at a slower rate (Rivkin, Bosch et al. 1986; Manahan, Davis et al. 1993; Feehan, Grauman-Boss et al. In Review). This implies that the pelagic period for larvae would lengthen greatly, and although microbe-induced mortality is reduced, the incidence of mortality by predation or offshore transport would be higher, assuming time-dependent relationship (Young and Chia 1987; Rumrill 1990).

Together, planktotrophic larvae in a phytoplankton bloom face two primary feeding environments: high particulate exogenous nutrients and high microbial load with pathogenic characteristics, or low particulate exogenous nutrients and low microbial load. To inhabit the former environment larvae would need to maintain an elevated metabolic rate (that promotes an accelerated development) as well as an elevated immune system (to combat the higher incident of pathogens). On the other hand, the latter requires the opposite, where low particulate exogenous nutrients and the likelihood of pathogens suppresses both metabolic activity and the need for an elevated immune system. Previous research using a diversity of animals has shown a well-characterized trade-off between the metabolic cost for growth, development, and body maintenance and functional capacity of the immune system to fend against pathogens (Lochmiller and Deerenberg 2000) (Figure 2).

Larvae in an environment with high food and a high microbial load are predicted to exhibit trade-offs between investing energetic input in growth, development, and body maintenance (*i.e.*, feeding) or the functional capacity of the immune system (*i.e.*, defend against pathogens), but not at maximum capacity for both. At present, research has, to our knowledge, yet to directly compare the trade-off of growth and immune function in marine invertebrate larvae, but a study with echinopleuti suggests this trade-off may occur (Carrier, King et al. 2015). Larvae of the sea urchin *Strongylocentrotus droebachiensis* were cultured in two different feeding environments: a high food reflective of the chlorophyll maximum and a low food which was phytoplankton-deprived, like surface and deeper water. Larvae in a chlorophyll maximum-like environment, as compared to diet-restricted individuals, exhibited higher levels of expression for genes involved with various metabolic processes. Genes involved in innate immunological response, on the other hand, were expressed at higher levels in larvae experiencing restricted diets, while other immune-associated genes were expressed at higher levels within the *ad libitum* condition (Figure 3). If these gene expression patterns persist in the natural environment, it would suggest that larvae concentrated in regions of high phytoplankton (and high microbial load) are unlikely to defend against environmental microbiota exhibiting pathogenic characteristics and, thus, are more susceptible to microbe-induced mortality when relying on the innate immune repertoire alone.

## Larval dysbiosis

Both host physiology (*e.g.*, innate immunity) and associated microbial flora act as the primary line of defense against pathogenic invaders. The composition and structure of this community, referred to as the microbiome, has been shaped over time in concert with host evolution and serves as an adaptive character for acclimating to environmental variation (Zilber-Rosenberg and Rosenberg 2008; Bordenstein and Theis 2015; Alberdi, Aizpurau et al. 2016; Carrier and Reitzel In Review). In the face of abiotic and/or biotic stressors, the composition of this community changes following the onset of a particular environmental change, which could be a stressor(s) (Carrier and Reitzel In Review). In certain cases, environment-mediated shifts in the composition of host-associated microbiota confer physiological acclimation (*e.g.*, recruitment of microbes for tolerance) while in other situations a shift in the microbiota promotes an increase in the abundance of pathogenic species. Moreover, the latter can occur in two primary fashions: microbiota previously associated with the host increase the expression of virulent gene products or microbiota initially not associated with the host dominate the microbial community. Both of these changes can result in the host entering a state of dysbiosis (Egan and Gardiner 2016).

Whether dysbiosis and/or pathogenic microbes is a significant cause of death in the plankton remains to be empirically measured in the field, but decades of larval culturing suggest it could be a significant contributing factor to mortality. First, to eliminate the environmental microbiota and to control microbial growth, cultures of benthic marine invertebrate larvae are often reared in 0.22-0.45 µm filtered seawater. Following the addition of exogenous nutrients to promote larval growth, incidences of unexplained larval mortality are sometimes observed, of which can be halted following the introduction of antibiotics (Strathmann 1987; Zhang, Chen et al. 2010); J Hodin, personal communication), suggesting a likely role for pathogenic microbes. Second, pathogens evading the immune system of planktotrophic larvae resulting in disease have been reported for ecologically important bivalves. For example, (Jeffries 1982) isolated three strains of *Vibrio* from diseased *Crassostrea gigas*, and determined that each strain can collapse cultures of *C. gigas* as well as *Ostra edulis* larvae within 48 h. The two examples above suggest marine bacterial pathogens can induce larval mortality, but may the environment (*i.e.*, feeding regime) mediate dysbiosis and subsequently mortality for marine invertebrate larvae?

A first attempt to determine if the feeding environment can influence larval microbial communities and dysbiosis suggest the answer may be yes. We reared larvae of the sea urchin *S. droebachiensis* in seawater containing the environmental microbial community (5 µm filtering) under exogenous nutrient conditions reflecting the chlorophyll maxima (fed *ad libitum*) and less biologically productive areas (10% *ad libitum* treatment) of the water column. As predicted, larvae reared in a feeding regime mirroring less biologically productive positions in the water column saw little to no mortality. On the other hand, larvae reared in a feeding regime reflective of the chlorophyll maxima exhibited high mortality over the course of development, such that most or all (~3,000) larvae died. In this latter culture, we sampled healthy (as indicated by active swimming and feeding) and dying (as indicated by exposed skeletal rods and degrading tissues, no swimming, and no feeding) larvae and assayed for their microbial communities (see caption of Figure 4 for methods).

In this experiment, healthy larvae associated with 42 operational taxonomic units (OTUs; species of bacteria) while dying larvae associate with 61 OTUs (Supplemental Figure 1), with four OTUs being found to only associate with healthy larvae, 23 OTUs to only associate with dying larvae, and 38 OTUs to associate with larvae in both states (Supplemental Figure 2). In focusing on associated OTUs representing ≥1% of the sequences (90.3% of healthy larvae data; 85.9% of dying larvae data), healthy larvae were dominated by *Vibrio* (60.4%) and *Flexibacter* (26.9%) while dying larvae associated most with *Vibrio* (25.0%) as well as nine OTUs between 4% and 13% (Figure 4). Most notably, in the transition from healthy to dying, *Vibrio* and *Flexibacter* decreased in abundance by 2.4-and 26.9-fold, respectively, while Colwelliaceae and *Thalassomonas* increased by 34.9-and 76.0-fold, respectively (Figure 4). Moreover, of the 14 OTUs associated with healthy and dying larvae (at ≥1% of the sequences), all exhibited ≥2.3-fold change in abundance, with most increasing coincident with disease and projected mortality (Figure 4). This shift in the composition of larval-associated microbiota in a feeding regime reflecting the chlorophyll maxima suggests that larvae may enter a state of dysbiosis, which subsequently leads to mortality or leave larvae more susceptible to pathogenic microbiota.

**Figure 1.**
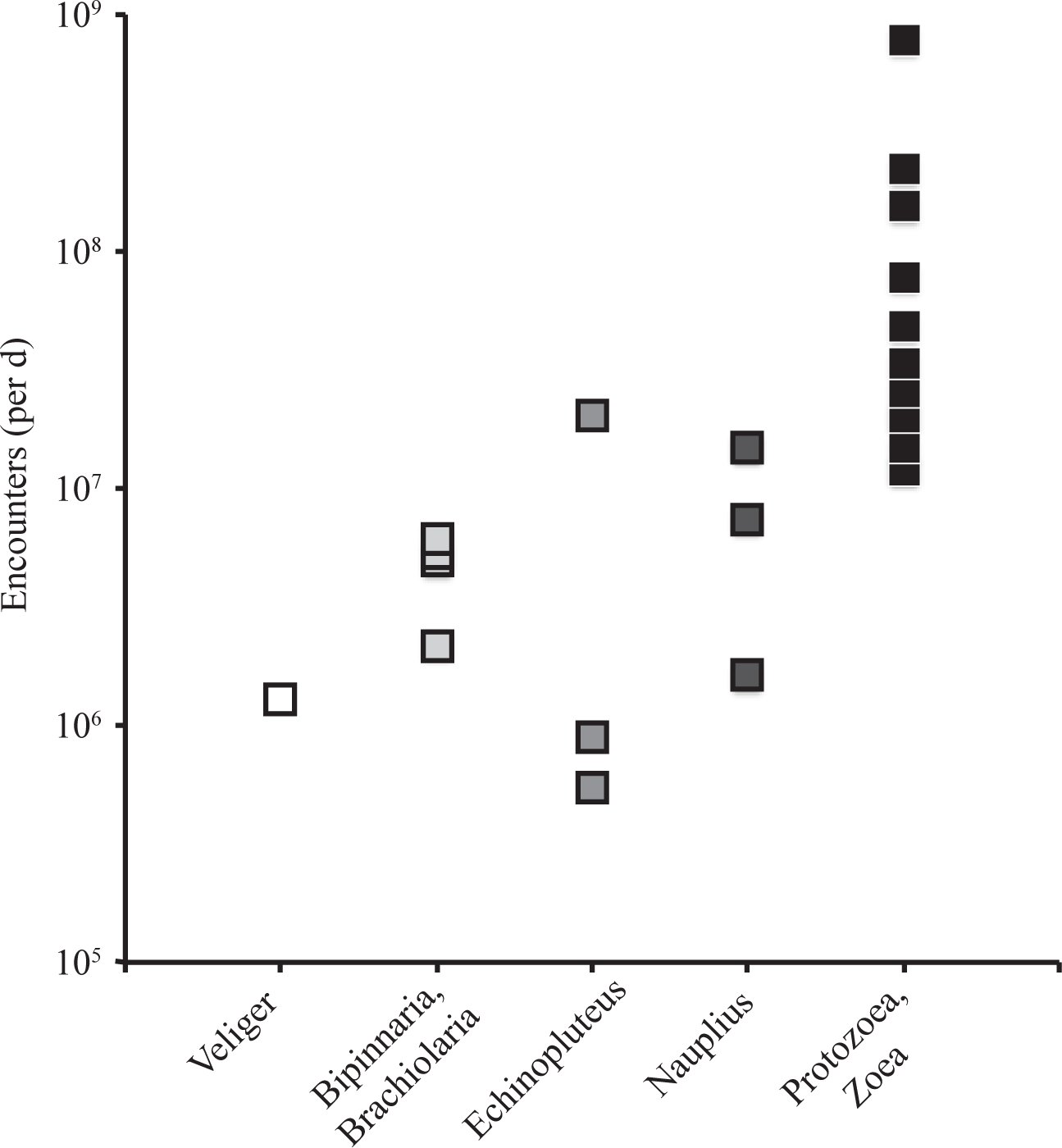
Predicted encounters between environmental microbiota and larval types by feeding. Predicted encounters were calculated as the product of maximum clearance rates of larvae reported by Strathmann (1987b) and mean bacterial abundance in the sea reported by Azam and Malfatti (2007). Here, each data point represents a species of benthic marine invertebrate orginally presented in Strathmann (1987b) that was reproduced and summarized in Supplemental Table 1.

**Figure 2.**
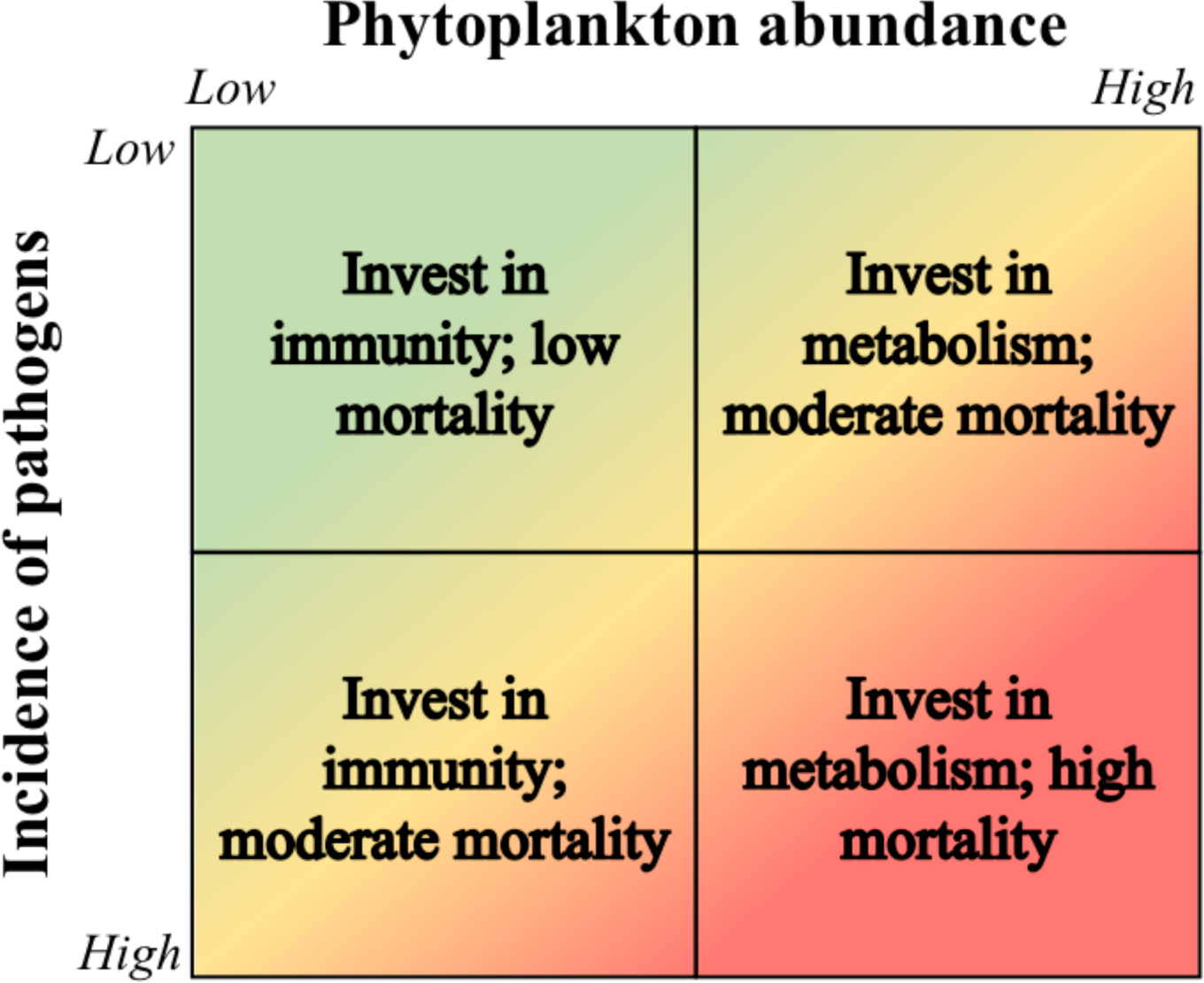
Metabolic-immune trade-off faced by larvae. Within the water column, larvae likely face two contrasting environments: high particulate exogenous nutrients and high microbial load with pathogenic characteristics, or low particulate exogenous nutrients and low microbial load. In the former environment, larvae would require an elevated metabolism for growth and development, and an elevated immunity to defend against pathogenic bacteria (bottom right). On the other hand, because both food and pathogens are lacking, the latter environment would require relatively much less investment in metabolism and immunity (top left). Because animals face a trade-off between investment in metabolism and immunity, the former environment is more likely to see incidences of mortality while the latter does not.

**Figure 3.**
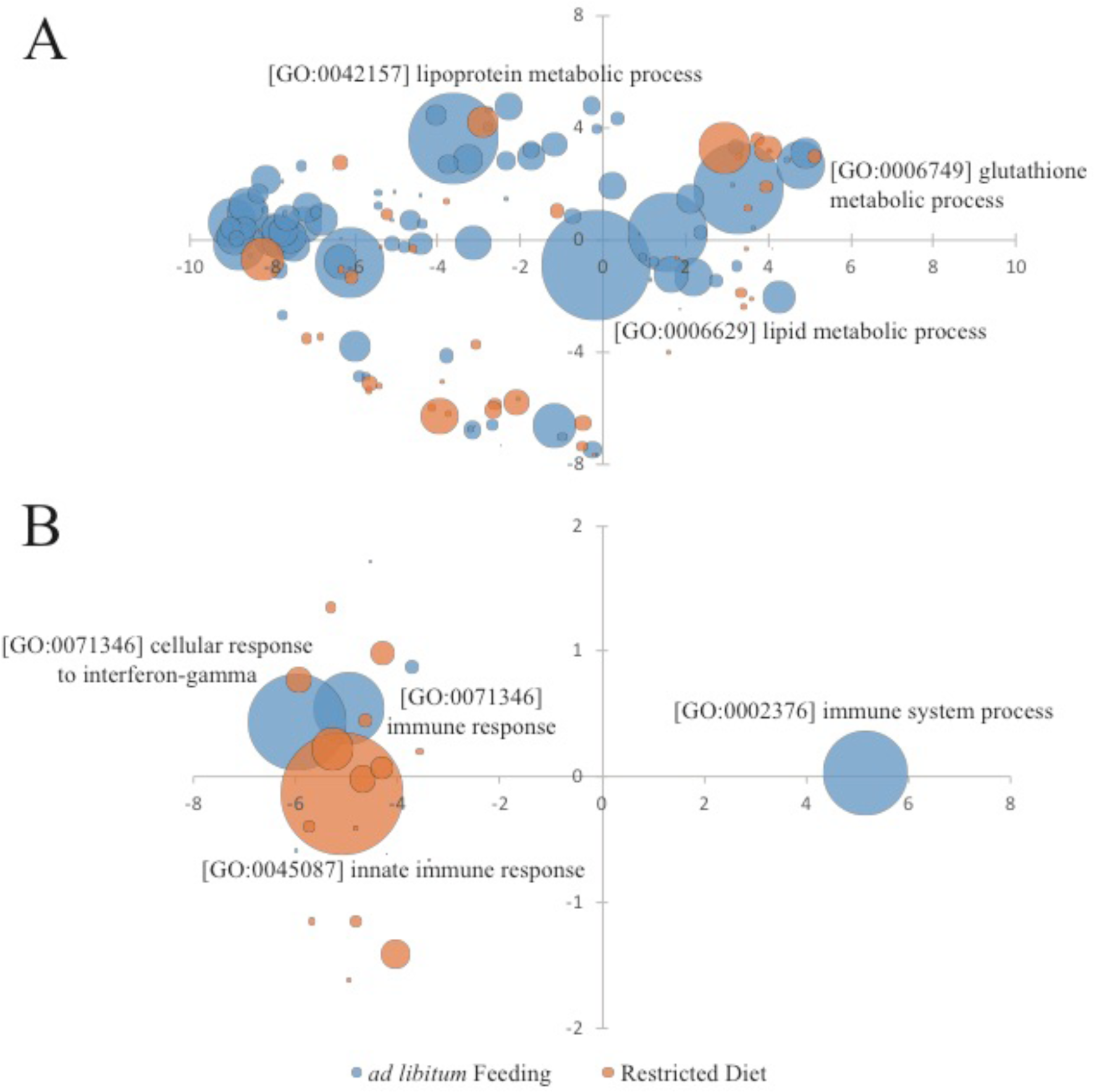
Differential gene expression and Gene Ontology (GO) analysis for *Strongylocentrotus droebachiensis* larvae in high nutrient (*ad libitum* feeding) and low nutrient (restricted diet) environments. Bubble size corresponds to differential in average GO expression (TPM) for each condition, with some of the more highly expressed GO groups labeled. REVIGO coordinates for GO groups correspond with semantic terminology with similar functional groups closer together (Supek, Bošnjak et al. 2011). (A) Transcripts associated with metabolism GO groups were significantly higher in larvae experiencing *ad libitum* feeding, with transcripts associated with diverse metabolic processes (lipid, lipoprotein, glutathione, etc.) having the highest expression. (B) Even though transcripts associated with the innate immune response were high for the restricted diet treatment, there were other immune related GO groups also expressed at high levels in *ad libitum* feeding environment. Although the experimental design for the data presented here (from Carrier et al. 2015) was not designed using the proposed hypothesis, the gene expression profiles of the host larvae follow similar predictive patterns as outlined by our hypothesis. For raw transcriptomic data see Supplemental File 1.

**Figure 4.**
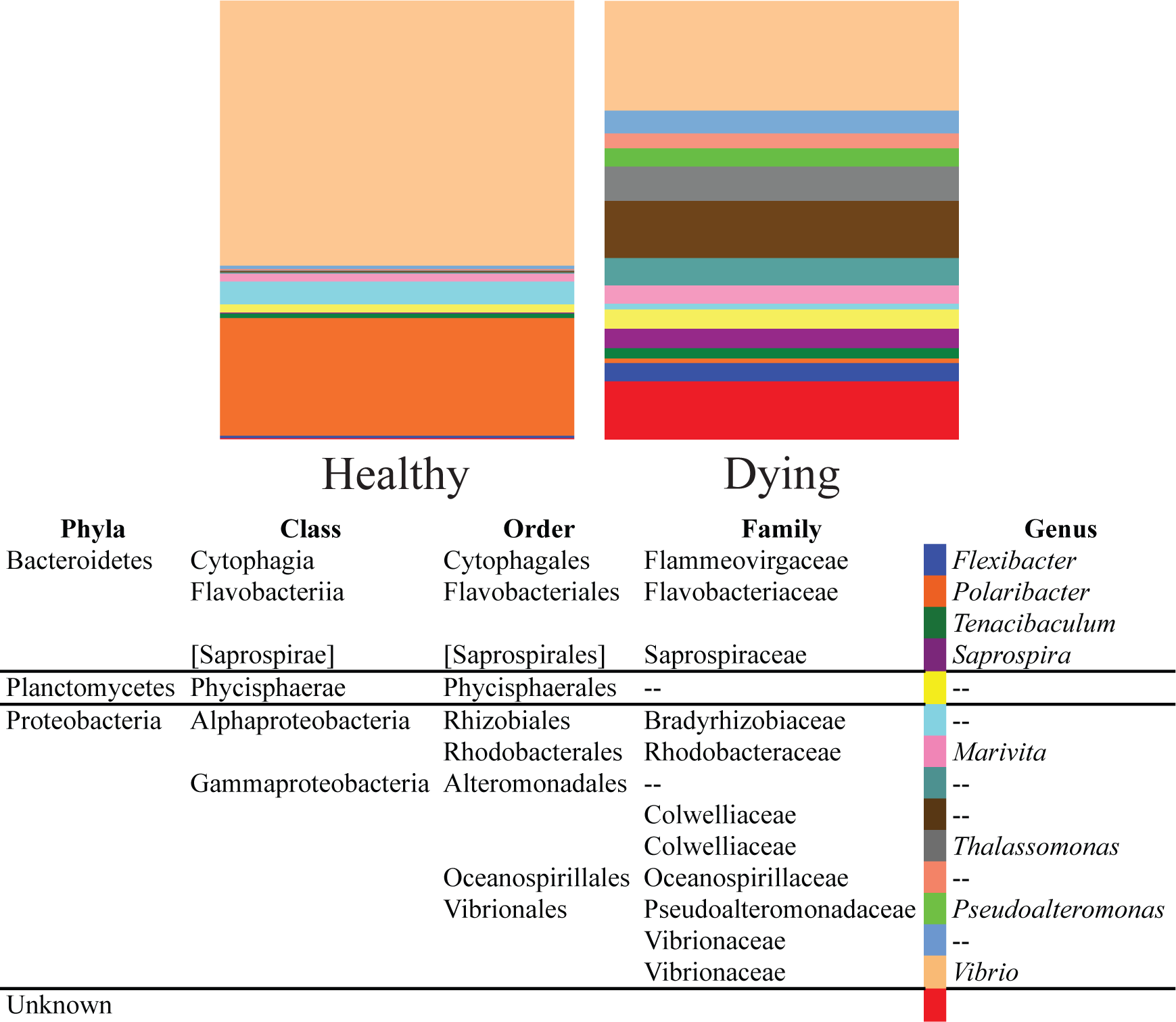
Bacterial communities associated with healthy and dying *Strongylocentrotus droebachiensis* larvae. Here, larvae of *Strongylocentrotus droebachiensis* larvae were fed *Rhodomonas ad libitum* in the presence of the environmental microbiota (5 µm filtered seawater), upon which high incidence of mortality was observed following two weeks of rearing. Both healthy and dying larvae were collected and stored in RNAlater. The larval hologenome was extracted and the V3/V4 region of 16S rRNA gene was amplified and sequenced. Forward and reverse raw read files were combined using PEAR (Zhang, Kobert et al. 2014), sequences in combined raw read files were trimmed using Trimmomatic (Bolger, Lohse et al. 2014), Fastq to Fasta using custom code, and taxonomically characterized using QIIME 1.9.1 (Caporaso, Kuczynski et al. 2010), with an OTU cutoff of 97%. Bar charts presented here, at the genus level, represent the microbiota associated with pools of approximately 50 healthy and dying *S. droebachiensis* larvae from the same culture.

## Experimental sampling

One of the most significant questions in the field of marine invertebrate life-history evolution is: why are reproductive modes used by species in their ecological niche? A common means to assess this question across the world’s oceans has been by correlating reproductive mode (*i.e.*, brooding, lecithotrophy, planktotrophy) with abiotic (temperature) and/or biotic (feeding regime) environment (Thorston 1950; Mileikovsky 1971; Marshall, Krug et al. 2012). Empirical studies of larvae in the natural environment are limited due to a number of factors, including their small size, their patchiness in the environment, and the large volume of water in which they may reside. However, in order to understand the intersection of larval biology, the feeding environment, and microbial communities, it is necessary to study these processes in the field. Laboratory studies would not accurately depict larvae physiology when feeding on complex food or the diversity of microbial assemblies because many species cannot be cultured in the laboratory (Carrier & Reitzel In Review). In addition, laboratory studies are not able to account for larval vertical position in the water column as this may also affect their odds of survival (Figure 1; Figure 2). Our understanding of the selective pressure pathogens and/or environment-mediated dysbiosis have on life-history evolution cannot broadly be assessed; however, proper experimental design would facilitate investigations of the contribution that the environmental microbiota and pathogens has on the planktotrophic life history.

Planktotrophic larvae develop from small, energy-poor eggs with only enough maternal input to complete embryogenesis and develop feeding structures, requiring an energetic contribution from exogenous resources to reach competency for metamorphosis. Planktotrophs require longer periods in the water column, primarily because of the relatively large quantity of energy needed to complete development, thus resulting in an extensive dispersal often 10s to 100s of kilometers (Thorston 1950; Mileikovsky 1971; Strathmann 1985; Shanks 2009). Many groups of benthic marine invertebrates primarily reproduce via planktotrophy due to the low energetic investment per offspring to facilitate a broad geographical distribution, increase gene flow, limit the likelihood of exposure to benthic predators and local extinction (Signor and Vermeij 1994; Pechenik 1999). These benefits are mostly offset by high rates of mortality, primarily from inadequate food conditions, temperature, offshore transport, and the inability to locate a suitable habitat for settlement (Thorston 1950; Young and Chia 1987; Rumrill 1990; Morgan 1995).

A source of mortality in the plankton not mentioned in seminal syntheses (Thorston 1950; Young and Chia 1987; Rumrill 1990; Morgan 1995) is microbe-induced mortality, whether directly by pathogen(s) or by dysbiosis. Our combined evaluation of the literature and empirical data suggest that feeding environment may mediate microbe-induced mortality in larvae through dysbiosis, and other studies suggest pathogen (*Vibrio*)-induced mortality. This is not to say, however, that microbes may not contribute to other sources of mortality in the plankton. For example, in the face of temperature stress larvae of the Great Barrier Reef sponge *Rhopaloeides odorabile* shift upwards of ~34% of their associated microbiota (Webster, Botte et al. 2011; Carrier and Reitzel In Review), but whether this results in *R. odorabile* larvae being more susceptible to pathogens or entering a state of dysbiosis remains unknown. This may be answered by rearing *R. odorabile* larvae (or larvae from other species) to elevated temperatures, exposing them to known larval pathogens, and assaying for mortality.

For a majority of species, we suggest ship-based field work would be necessary for environmentally relevant studies of larvae-microbes interactions (Figure 5). The physical, chemical, biological, and microbial parameters of the water column are characterized (*e.g.*, CTD), and larvae are subsequently sampled at the chlorophyll maximum as well as in less productive surface and bottom waters (Figure 5). To best preserve the larval hologenome, samples would be preserved (*e.g.*, RNAlater) immediately at sea, then the genomic DNA is extracted, 16S rRNA gene is amplified and sequenced (*e.g.*, MiSeq and 454-pyrosequencing), and microbiota are classified using bioinformatic programs (*e.g.*, QIIME and mothur) (Figure 5) (Williams and Carrier 2017).

**Figure 5.**
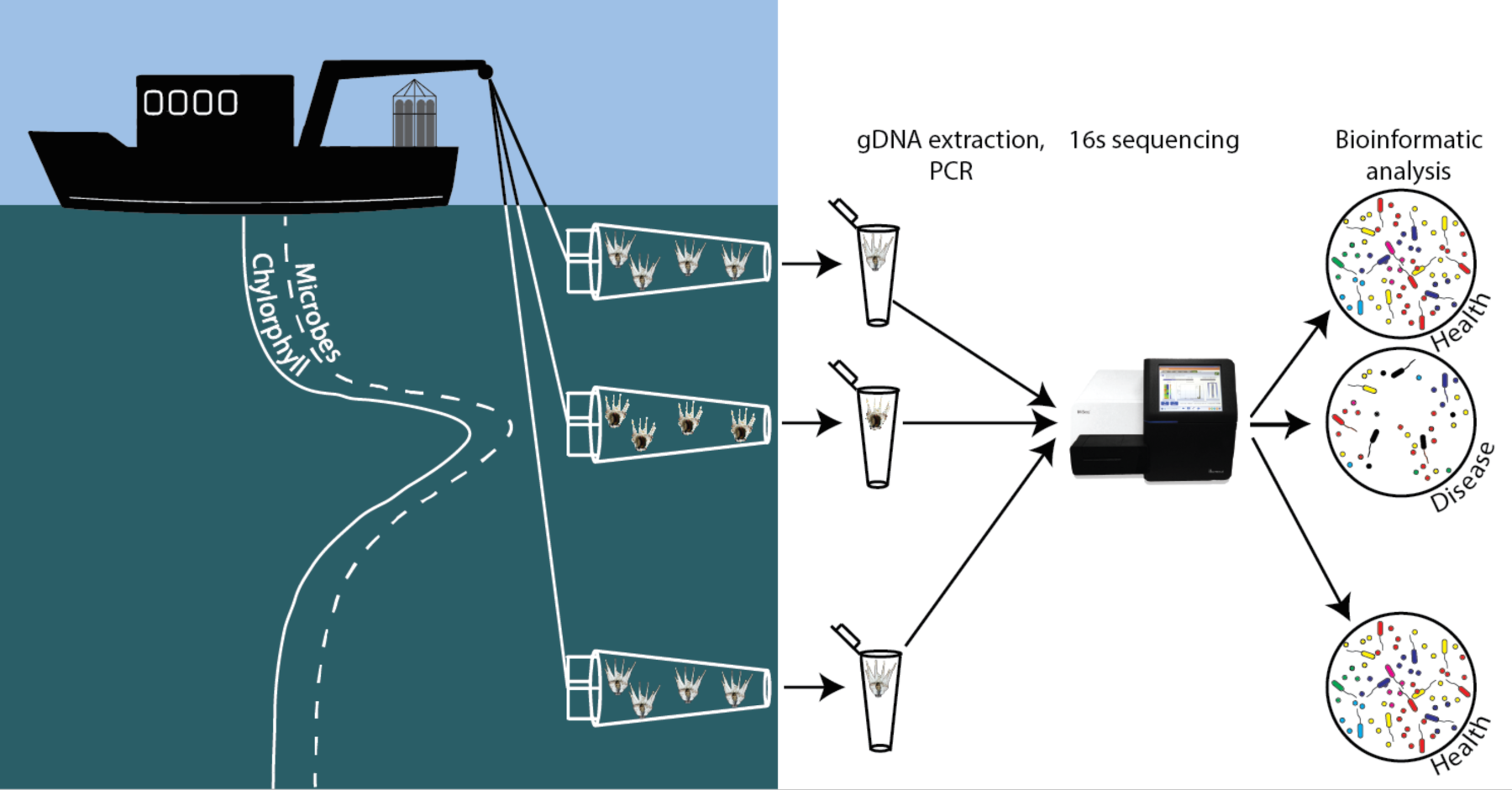
Experimental approach to characterize the intersection and dynamics of the environmental metagenome and larval microbiome. The hypothesis proposed here suggests that a life-history trade-off exists between feeding environment (and, therefore, larval growth and time spent in the plankton) and exposure to virulent microbes. Being that the microbial load, with particular species being more virulent in that setting, is elevated in areas of high biological productivity and thus for larval growth, avoidance of microbial-mediated diseases is higher in areas of reduced productivity (*i.e.*, surface or deep waters). As such, to avoid microhabitats with elevated microbial load and subsequent virulence, larvae may have adopted different mechanisms to cope with this trade off, including behavioral avoidance, changes in expression of the immune system, or shifts from dependence on exogenous food. Empirically testing these hypotheses requires an interdisciplinary approach that combines profiling the water column for oceanographic properties (*i.e.*, CTD), sampling larvae in different portions of the water column (*i.e.*, plankton tows), identifying and comparing the microbial community with metagenomics (*e.g.*, Illumina MiSeq, QIIME) and experimentally testing larval performance and survival in laboratory cultures. This type of approach would test whether larvae in areas of high chlorophyll exhibit dysbiosis and a subsequent microbial-mediated mortality.

This proposed fieldwork would aim to determine if vertical position in the water column and relative feeding regime would be an important initial step to determine whether oceanographic features of the sea influence the biology and ecology of marine invertebrate larvae and their associated microbiota, and further how this then contributes to mortality in the plankton. Going forward, it remains paramount to compliment current endeavors in larval ecology and life-history evolution with the study of associated microbiota towards a holistic effort to understand the evolutionary ecology of benthic marine invertebrates and their larvae. Approaches that address these questions may be best served at the intersection of oceanography and life sciences (Theis, Dheilly et al. 2016; Carrier and Reitzel In Review).

## Acknowledgements

We thank Colette Feehan and Richard Strathmann for provoking discussions, and members of the Reitzel Lab and Richard Strathmann for providing comments on an earlier version of this manuscript. This work was funded by the National Science Foundation (TJC, AMR), the National Institute of Health (AMR), the Human Frontiers Science Program (AMR), North Carolina Sea Grant (TJC, AMR), the Charles Lambert Memorial Endowment at the Friday Harbor Laboratories (TJC), and Sigma Xi (TJC).

---

**Figure S1.**
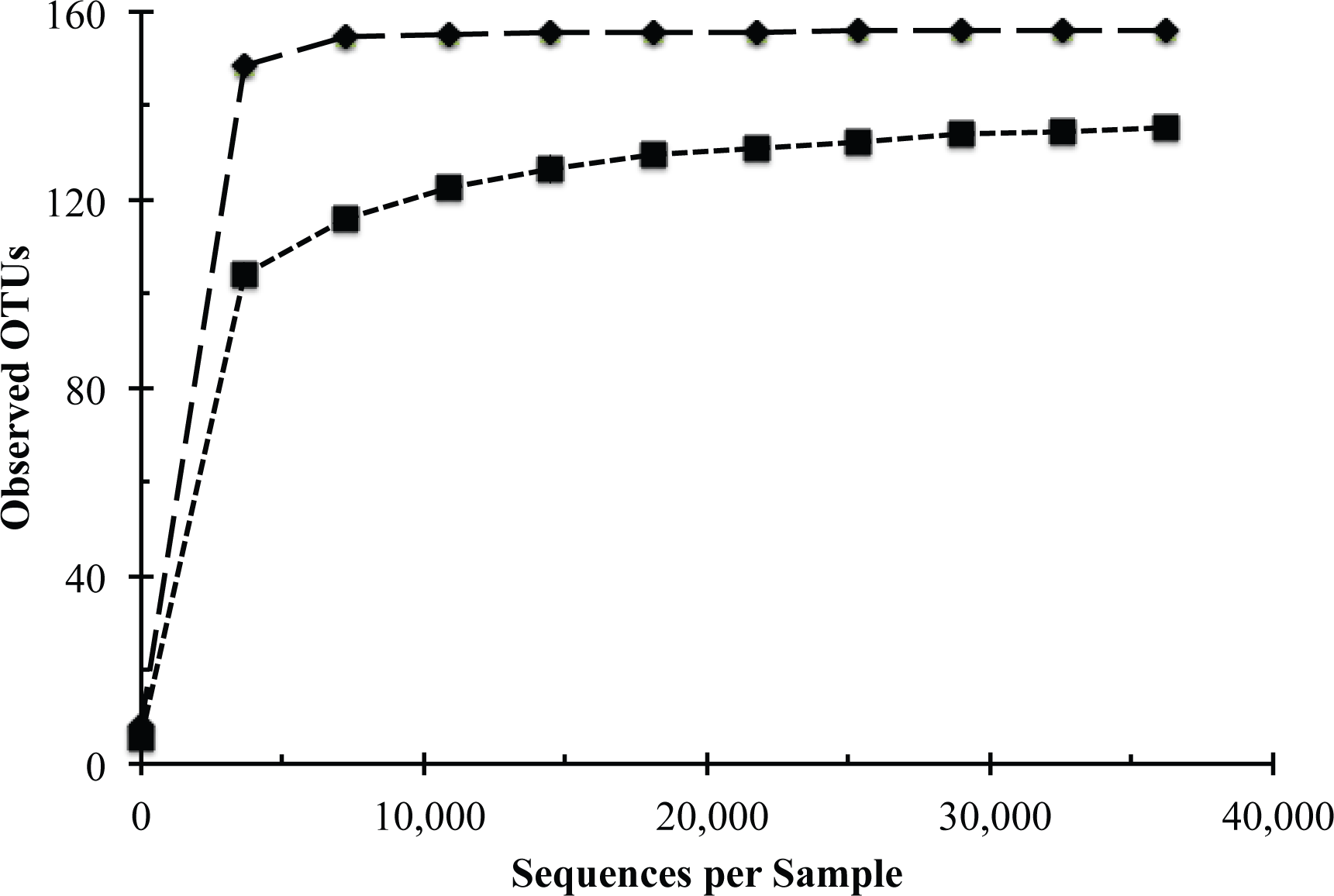
Alpha rarefaction of observed number of operational taxonomic units (OTUs) in each larval sample with diamonds representing dying larvae and squares representing healthy larvae. This analysis was completed using the alpha_rarefaction.py script in QIIME 1.9.1 (Caporaso, Kuczynski et al. 2010), with the rarefaction curves being recreated in Adobe Illustrator CS6.

**Figure S2.**
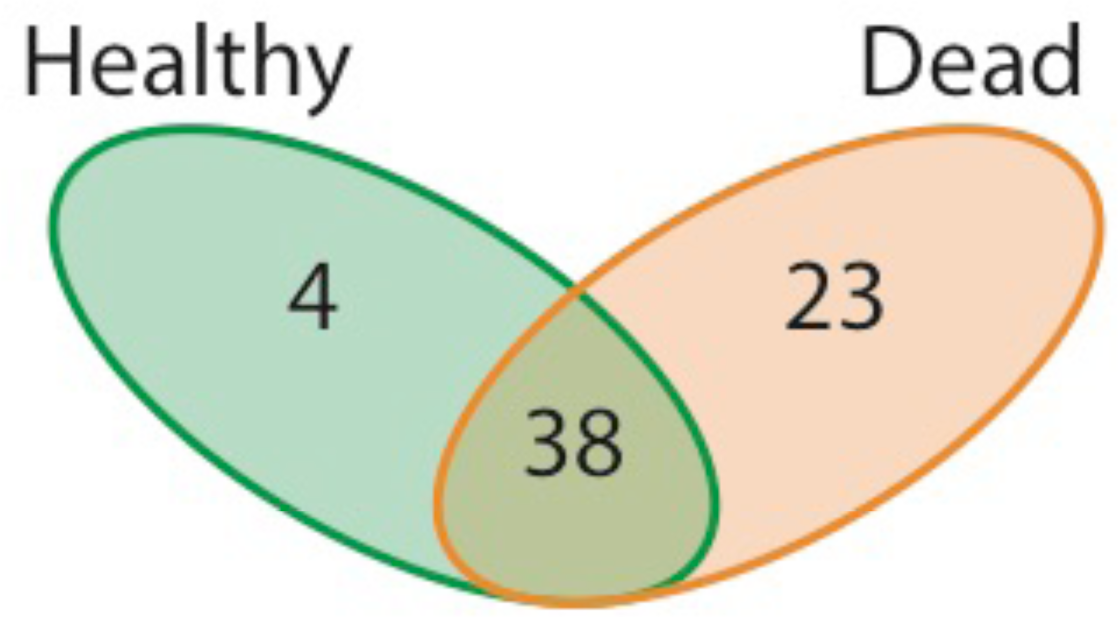
Shared and unique host-associated microbiota of healthy and dying *Strongylocentrotus droebachiensis* larvae. Healthy *S. droebachiensis* larvae associated with 42 operational taxonomic units (OTUs) while dying larvae associate with 61 OTUs, with four OTUs being found to only associate with healthy larvae, 23 OTUs to only associate with dying larvae, and 38 OTUs to associate with larvae in both states. OTU summaries were generated using the summarize_taxa_through_plots.py script in QIIME 1.9.1 (Caporaso, Kuczynski et al. 2010) and the shared and unique OTUs were then counted by hand. This Venn diagram summary was created using Adobe Illustrator CS6.

| <u>Species</u> | <u>Stage</u> | <u>Clearance (mL/d)</u> | <u>Encounters (per d)</u> |
| --- | --- | --- | --- |
| **Veliger** |  |  |  |
| *Mytilus edulis* | | 1.3 | $1.3 \times 10^6$ |
| *Crassostrea gigas* | | 0.1 | $1.3 \times 10^5$ |
| *Ostrea edulis* | | 0.9 | $9.0 \times 10^5$ |
| *Teredo navalis* | | 4.1 | $4.1 \times 10^6$ |
| *Illyanassa obsoleta* | | 1.8 | $1.8 \times 10^6$ |
| **Bipinnaria, Brachiolaria** |  |  |  |
| *Acanthaster planci* | | 5.0 | $5.0 \times 10^6$ |
| *Luidia foliolata* | | 6.3 | $6.3 \times 10^6$ |
| *Evasterias troscheli* | | 2.2 | $2.2 \times 10^6$ |
| *Pisaster ochraceus* | | 5.3 | $5.3 \times 10^6$ |
| **Echinopluteus** |  |  |  |
| *Paracentrotus lividus* | | 0.9 | $9.0 \times 10^5$ |
| *Arbacia lixula* | | 0.5 | $5.4 \times 10^5$ |
| *Dendraster excentricus* | | 20.2 | $2.0 \times 10^7$ |
| **Nauplius** |  |  |  |
| *Calacus pacificus* | N3 | 1.6 | $1.6 \times 10^6$ |
| | N4 | 7.5 | $7.5 \times 10^6$ |
| | N5 | 15.0 | $1.5 \times 10^7$ |
| | N6 | 15.0 | $1.5 \times 10^7$ |
| **Protozoa, Zoea** |  |  |  |
| *Penaeus indicus* | P1 | 12.0 | $1.2 \times 10^7$ |
| | P2 | 12.0 | $1.2 \times 10^7$ |
| | P3 | 26.0 | $2.6 \times 10^7$ |
| | Z1 | 34.0 | $3.4 \times 10^7$ |
| | Z2 | 20.0 | $2.0 \times 10^7$ |
| | Z3 | 20.0 | $2.0 \times 10^7$ |
| *Sergestes similis* | P1 | 14.7 | $1.5 \times 10^7$ |
| | P2 | 49.0 | $4.9 \times 10^7$ |
| | P3 | 157.0 | $1.6 \times 10^8$ |
| | P3 | 79.0 | $7.9 \times 10^7$ |
| | Z2 | 800.0 | $8.0 \times 10^8$ |
| *Chionoecetes bairdi* | Z2 | 221.5 | $2.2 \times 10^8$ |

## References

Alberdi, A., O. Aizpurau, et al. (2016) Do vertebrate gut metagenomes confer rapid ecological adaptation. Trends in Ecology & Evolution, 311(9), 689–699.

Arellano, S.M., A.M. Reitzel, et al. (2012) Variation in vertical distribution of sand dollar larvae relative to haloclines, food, and fish cues. Journal of Experimental Marine Biology and Ecology, 414-415, 28–37.

Azam, F. (1998) Microbial control of oceanic carbon flux: the plot thickens. Science, 280(5364), 694–696.

Azam, F., F. Malfatti (2007) Microbial structuring of marine ecosystems. Nature Reviews Microbiology, 5, 782–791.

Bolger, A., M. Lohse, et al. (2014) Trimmomatic: a flexible trimmer for Illumina sequence data. Bioinformatics, 30, 2114–2120.

Bordenstein, S.R., K.R. Theis (2015) Host Biology in Light of the Microbiome: Ten Principles of Holobionts and Hologenomes. PLoS Biology, 13(8), e1002226.

Brum, J.R., J.C. Ignacio-Espinoza, et al. (2015) Patterns and ecological drivers of ocean viral communities. Science, 348(6237), 1261498.

Burgess, S.C., M.L. Baskett, et al. (2015) When is dispersal for dispersal? Unifying marine and terrestrial perspectives. Biological Reviews, 91(3), 867–882.

Caporaso, J.G., J. Kuczynski, et al. (2010) QIIME allows analysis of high-throughput community sequencing data. Nature Methods, 7(5), 335–336.

Carrier, T.J., B.L. King, et al. (2015) Gene expression changes associated with the developmental plasticity of sea urchin larvae in response to food availability. Biological Bulletin, 228, 171–180.

Carrier, T.J., A.M. Reitzel (In Review) The hologenome across environments and the implications of a host-associated microbial repertoire.

Carrier, T.J., A.M. Reitzel, et al. (2017). Evolutionary Ecology of Marine Invertebrate Larvae, Oxford University Press

Dekshenieks, M.M., P.L. Donaghay, et al. (2001) Temporal and spatial occurance of thin phytoplankton layers in relation to physical processess. Marine Ecology Progress Series, 223, 61–71.

Egan, S., M. Gardiner (2016) Microbial dysbiosis: rethinking disease in marine ecosystems. Frontiers in Microbiology, 7, 991.

Feehan, C.J., B.C. Grauman-Boss, et al. (In Review) Kelp detritus provides high-quality food for sea urchin larvae.

Galac, M.R., I. Bosch, et al. (2016) Bacterial communities of oceanic sea star (Asteroidea: Echinodermata) larvae. Marine Biology, 163, 162.

Gallagher, S.M., J.B. Waterbury, et al. (1994) Efficient grazing and utilization of the marine cyanobacterium Synechococcus sp. by larvae of the bivalve Mercenaria mercenaria. Marine Biology, 119(2), 251–259.

Gilbert, S.F., T.C.G. Bosch, et al. (2015) Eco-Evo-Devo: developmental symbiosis and developmental plasticity as evolutionary agents. Nature Reviews Genetics, 16(10), 611–622.

Hart, M.W., R.R. Strathmann (1994) Functional consequences of phenotypic plasticity in echinoid larvae. Biological Bulletin, 186(3), 291–299.

Harvell, C.D., C.E. Mitchell, et al. (2002) Climate warming and disease risks for terrestrial and marine biota. Science, 296, 2158–2162.

Hodin, J., M.C. Ferner, et al. (2017) I feel that! Fluid dynamics and sensory aspects of larval settlement across scales. In: T.J. Carrier, A.M. Reitzel & A. Heyland (Eds). Evolutionary Ecology of Marine Invertebrate Larvae. Oxford University Press, Oxford, UK.

Jaeckle, W.B. (2017) Physiology of larval feeding In: T.J., Carrier A.M. Reitzel & A. Heyland (Eds). Evolutionary Ecology of Marine Invertebrate Larvae. Oxford University Press, Oxford, UK.

Jeffries, V.E. (1982) Three vibrio strains pathogenic to larvae of Crassostrea gigas and Ostrea edulis. Aquaculture, 29, 201–226.

Lochmiller, R.L., C. Deerenberg (2000) Trade-offs in evolutionary immunology: just what is the cost of immunity? Oikos, 88, 87–98.

Manahan, D.T., J.P. Davis, et al. (1993) Bacteria-free sea urchin larvae: selective uptake of neutral amino acids from seawater. Science, 220(4593), 204–206.

Marshall, D.J., P.J. Krug, et al. (2012) The biogeography of marine invertebrate life histories Annual Review of Ecology, Evolution, and Systematics 43, 97–114.

McAlister, J.S., B.G. Miner (2017) Phenotypic plasticity of feeding structures in marine invertebrate larvae. In: T.J. Carrier A.M. Reitzel & A. Heyland (Eds). Evolutionary Ecology of Marine Invertebrate Larvae. Oxford University Press, Oxford, UK.

McEdward, L.R. (1984) Morphometric and metabolic analysis of the growth and form of an echinopluteus. Journal of Experimental Marine Biology and Ecology, 82, 259–287.

McFall-Ngai, M., M.G. Hadfield, et al. (2013) Animals in a bacterial world, a new imperative for the life sciences. Proceedings of the National Academy of Sciences of the United States of America, 110(9), 3229–3236.

McFall-Ngai, M., E.G. Ruby (2000) Developmental biology and marine invertebrate symbioses. Current Opinion in Microbiology, 3, 603–607.

McKenny, D., D.G. Allison (1995) Effects of growth rate and nutrient limitation on virulence factor production in Burkholderia cepacia. Journal of Bacteriology, 177(14), 4140–4143.

Metaxas, A., L.S. Mullineaux, et al. (2009) Distribution of echinoderm larvae relative to the halocline of a salt wedge. Marine Ecology Progress Series, 377, 157–168.

Mileikovsky, S.A. (1971) Types of larval development in marine bottom invertebrates, their distribution and ecological significance: a re-evalution. Marine Biology, 10, 193–213.

Morgan, S.G. (1995) Life and death in the plankton: larval mortality and adaptation. In: L. McEdwards (Ed). Ecology of Marine Invertebrate Larvae. CRC Press: 279–321.

Needham, D.M., J.A. Fuhrman (2016) Pronounced daily succession of phytoplankton, archaea and bacteria following a spring bloom. Nature Microbiology, 1, 16005.

Pechenik, J.A. (1999) On the advantages and disadvantages of larval stages in benthic marine invertebrate life cycles. Marine Ecology Progress Series, 177, 269–297.

Pespeni, M.H., E. Sanford, et al. (2013) Evolutionary change during experimental ocean acidification. Proceedings of the National Academy of Sciences of the United States of America, 110(17), 6937–6942.

Rivkin, R.B., I. Bosch, et al. (1986) Bacterivory: a novel feeding mode for asteroid larvae. Science, 233, 1311–1314.

Rosenberg, E., G. Sharon, et al. (2009) The hologenome theory of evolution contains Lamarckian aspects within a Darwinian framework. Environmental Microbiology, 11(12), 2959–2962.

Rumrill, S.S. (1990) Natural mortality of marine invertebrate larvae. Ophelia, 32, 163–198.

Shanks, A.L. (2009) Pelagic larval duration and dispersal distance revisited. Biological Bulletin, 216, 373–385.

Signor, P.W., G.J. Vermeij (1994) The plankton and the benthos: origins and early history of an evolving relationship. Paleobiology, 20, 297–319.

Starr, M., J. Himmelman, et al. (1990) Direct coupling of marine invertebrate spawning with phytoplankton blooms. Science, 247, 1071–1074.

Strathmann, M.F. (1987). Reproduction and development of marine invertebrates of the northern Pacific coast: data and methods for the study of eggs, embryos, and larvae. University of Washington Press.

Strathmann, R.R. (1985) Feeding and nonfeeding larval development and life-history evolution in marine invertebrates. Annual Review of Ecology and Systematics, 16, 339–361.

Strathmann, R.R. (1990) Why life histories evolove different in the sea. American Zoologist, 30(1), 197–207.

Strathmann, R.R., D. Grunbaum (2006) Good eaters, poor swimmers: compromises in larval form. Inegrative and Comparative Biology, 46, 312–322.

Sunagawa, S., L.P. Coelho, et al. (2015) Structure and function of the global ocean microbiome. Science, 348(6237), 1261359.

Supek, F., M. Bošnjak, et al. (2011) REVIGO summarizes and visualizes long lists of gene ontology terms. PLoS ONE, 6(7), e21800.

Theis, K.R., N.M. Dheilly, et al. (2016) Getting the hologenome concept right: An eco-evolutionary framework for hosts and their microbiomes. mSystems, 1(2), e00028–00016.

Thorston, G. (1950) Reproductive and larval ecology of marine bottom invertebrates. Biological Reviews, 25, 1–45.

Webster, N.S., E.S. Botte, et al. (2011) The larval sponge holobiont exhibits high thermal tolerance. Environmental Microbiology Reports, 3(6), 756–762.

Williams, E.A., T.J. Carrier (2017) An ‘-omics perspective on marine invertebrate larvae. In: T.J. Carrier, A.M. Reitzel & A. Heyland (Eds). Evolutionary Ecology of Marine Invertebrate Larvae. Oxford University Press, Oxford, UK.

Young, C.M., F.-S. Chia (1987) Abundance and distribution of pelagic larvae as influenced by predation, behavior, and hydrographic factors. In: A.C. Giese, J.S. Pearse & V.B. Pearse (Eds). Reproduction of Marine Invertebrates IX. General aspects: seeking unity in diveristy. Blackwell Scienfic Publications, Palo Alto, California.

Zhang, C., G. Chen, et al. (2010) Etiology of rotting edges syndrome in cultured larval Apostichopus japonicus at auricularia stage and analysis of reservoir of pathogens. Acta Microbiologica Sinica 49(5), 631–637.

Zhang, J., K. Kobert, et al. (2014) PEAR: a fast and accurate Illumina Paired-End reAd mergeR. Bioinformatics, 30, 614–620.

Zilber-Rosenberg, I., E. Rosenberg (2008) Role of microorganisms in the evolution of animals and plants: the hologenome theory of evolution. FEMS Microbiology Reviews, 32(5), 723–735.

